# L1 and B1 repeats blueprint the spatial organization of chromatin

**DOI:** 10.1101/802173

**Authors:** J. Yuyang Lu, Lei Chang, Tong Li, Ting Wang, Yafei Yin, Ge Zhan, Ke Zhang, Michelle Percharde, Liang Wang, Qi Peng, Pixi Yan, Hui Zhang, Xue Han, Xianju Bi, Wen Shao, Yantao Hong, Zhongyang Wu, Peizhe Wang, Wenzhi Li, Jing Zhang, Zai Chang, Yingping Hou, Pilong Li, Miguel Ramalho-Santos, Jie Na, Wei Xie, Yujie Sun, Xiaohua Shen

## Abstract

Despite extensive mapping of three-dimensional (3D) chromatin structures, the basic principles underlying genome folding remain unknown. Here, we report a fundamental role for L1 and B1 retrotransposons in shaping the macroscopic 3D genome structure. Homotypic clustering of B1 and L1 repeats in the nuclear interior or at the nuclear and nucleolar peripheries, respectively, segregates the genome into mutually exclusive nuclear compartments. This spatial segregation of L1 and B1 is conserved in mouse and human cells, and occurs dynamically during establishment of the 3D chromatin structure in early embryogenesis and the cell cycle. Depletion of L1 transcripts drastically disrupts the spatial distributions of L1- and B1-rich compartments. L1 transcripts are strongly associated with L1 DNA sequences and induce phase separation of the heterochromatin protein HP1α. Our results suggest that genomic repeats act as the blueprint of chromatin macrostructure, thus explaining the conserved higher-order structure of chromatin across mammalian cells.

## INTRODUCTION

The mammalian genome is folded nearly one million-fold within the nucleus of a cell (Annunziato, 2008). Hierarchical organization of genome folding at different scales is central to gene expression, cellular function and development, and alterations in genome organization often lead to disease (Dekker et al., 2017; Zane et al., 2014). Microscopic and 3C-based approaches reveal that chromatin forms a “fractal globule” with compartments, domains, and loop interactions (Boettiger et al., 2016; Dekker et al., 2002; Mirny, 2011; Wang et al., 2016). At the megabase scale, chromatin is subdivided into two spatially segregated compartments, arbitrarily labeled as A and B, with distinct transcriptional activity (Lieberman-Aiden et al., 2009). The A and B compartments also differ in other features, including gene density, CpG frequency, histone modifications, and DNA replication timing (Bonev and Cavalli, 2016; Dekker et al., 2013; Gibcus and Dekker, 2013; Rivera-Mulia and Gilbert, 2016). Simulation of three-dimensional (3D) genome structures predicted that the euchromatic A compartment adopts a central position, whereas the heterochromatic B compartment moves towards the nuclear periphery and perinucleolar regions (Buchwalter et al., 2019; Stevens et al., 2017). Within compartments at the kilobase-to-megabase scale, chromatin is organized in topologically associated domains (TADs), which serve as functional platforms for physical interactions between co-regulated genes and regulatory elements (Dixon et al., 2012). At a finer scale, TADs are divided into smaller loop domains, in which distal regulatory elements such as enhancers come into direct contact with their target genes via chromatin loops (Kadauke and Blobel, 2009).

The organization of the nuclear genome into euchromatin and heterochromatin, which closely match the A and B compartments, respectively, appears to be conserved from ciliates to humans and has been maintained in eukaryotes over 500 million years of evolution (Solovei et al., 2016). Intriguingly, most A/B compartments and TADs seem to be invariant in different mouse and human cell types or during cell-fate transition, whereas sub-TAD loops are more variable to facilitate differential gene expression (Dixon et al., 2015; Stadhouders et al., 2018). The insulator protein CTCF (CCCTC-binding factor) and the ring-shaped cohesin complex, considered as the weavers of genome organization, co-localize at the boundaries and anchor regions of contact domains and loops (Rao et al., 2014; Splinter et al., 2006). However, depletion of CTCF or cohesin disrupts some sub-TAD loop contacts, but fails to affect the formation and maintenance of A/B compartments in cells (Bintu et al., 2018; Gassler et al., 2017; Haarhuis et al., 2017; Nora et al., 2017; Nuebler et al., 2018; Rao et al., 2017; Schwarzer et al., 2017; Wutz et al., 2017). The direct molecular drivers, other than CTCF and cohesin, that govern compartmental domains are yet to be defined. A few mechanisms have been proposed for the formation of nuclear compartments, such as anchoring heterochromatin to the nuclear lamina (Falk et al., 2019; Guelen et al., 2008; van Steensel and Belmont, 2017; Wijchers et al., 2015; Zullo et al., 2012), preferential attraction of chromatin harboring similar histone modifications and regulators (Jost et al., 2014; Strom et al., 2017; Wang et al., 2016), and hypothetical models involving pairing of homologous sequences mediated by active transcription or phase separation (Cook, 2002; Cook and Marenduzzo, 2018; Rowley et al., 2017; Tang, 2017; van de Werken et al., 2017). A recent study based on imaging and polymer simulations proposes that interactions between heterochromatic regions are crucial for compartmentalization of the genome in inverted and conventional nuclei (Falk et al., 2019). However, the molecular basis that drives heterochromatin compartmentalization and segregation from active domain remains unclear (Bouwman and de Laat, 2015; Pueschel et al., 2016).

Repetitive elements comprise about half of human and mouse genomes (Biemont, 2010). Once regarded as genomic parasites (Orgel and Crick, 1980), retrotransposons have been recently implicated in playing active roles in re-wiring the genome and gene expression programs in diverse biological processes (Bourque et al., 2008; Chuong et al., 2016; Durruthy-Durruthy et al., 2016; Grow et al., 2015; Liu et al., 2018; Lynch et al., 2011; Rebollo et al., 2012; Wang et al., 2014). Repetitive DNA is speculated to have a role in nuclear organization (Falk et al., 2019; Solovei et al., 2016; Tang, 2017). Considering the diverse nature and widespread distribution of repeats in mammalian genomes (Sundaram and Wang, 2018), it is imperative to determine whether and how repetitive sequences are involved in this process. Long and short interspersed nuclear elements (LINEs and SINEs, respectively) are the two predominant subfamilies of retrotransposons in most mammals (Mandal and Kazazian, 2008). L1 (also named as LINE1 or LINE-1) is the most abundant subclass of LINEs, and also constitutes the most abundant type of all repeat subclasses, making up to 18.8% and 16.9% (0.9∼1.0 million copies) of the genome in mouse and human, respectively (Taylor et al., 2013). B1 in mouse and its closely related, primate-specific Alu elements in human are the most abundant subclass of SINEs, constituting 2.7%∼10.6% (0.6∼1.3 million copies) of mouse and human genomes (Lander et al., 2001; Mouse Genome Sequencing et al., 2002). L1 and B1/Alu have distinct nucleotide compositions and sequence lengths. While Alu elements are ∼300 bp long and rich in G and C nucleotides, L1 elements are 6–7 kb long and AT-rich (Jurka et al., 2004). A previous study of metaphase chromosome banding showed roughly inverse distributions of L1 and Alu elements in chromosomal regions with distinct biochemical properties (Korenberg and Rykowski, 1988). Further studies reported that Alu/B1 elements are preferentially enriched in regions that are generally gene-rich, whereas L1 elements are enriched in gene-poor regions, which tend to be located in lamina-associated heterochromatin (Deininger, 2011; Meuleman et al., 2013; Wijchers et al., 2015). However, systematic mapping of L1 and B1 elements in the genome and the nuclear space is still lacking.

Here, we report that L1 and B1/Alu repeats harbor the basic structural information in their DNA sequences to instruct the 3D organization of mammalian genomes. Co-segregation of B1/Alu and L1 repeats with the A and B compartments, respectively, is highly conserved in different mouse and human cell types. B1/Alu-rich regions cluster in the interior of the nucleus, while L1-rich regions cluster at the nuclear and nucleolar periphery. This pattern of homotypic clustering and segregation is dynamically established in the cell cycle and in early embryogenesis. L1 RNA is critically required for the formation and maintenance of compartment segregation and the higher-order chromatin structure. Most likely, L1 RNA transacts the structural information in L1 DNA repeats by acting upon its own DNA sequences to promote phase separation of HP1α, thereby inducing heterochromatin formation. Our findings uncover a fundamental principle of 3D genome organization that is rooted in genomic DNA sequences, and also provide an answer to the perplexing phenomenon of the conservation and robustness of chromatin compartmentalization across mammalian species.

## RESULTS

### Homotypic clustering of B1 and L1 repeats segregates the A/B compartments

We analyzed the genomic positions of the major repeat subfamilies and observed positive correlations within the L1 and SINE B1 subfamilies, but strong inverse correlations between them (Figure S1A). This observation suggests that L1 and SINE B1 elements tend to be positioned away from each other in the genome, while repeats from the same subfamily tend to be clustered. The non-random positioning of repeat sequences in the genome prompted us to examine their relative distributions in the A/B compartments. Analysis of Hi-C data in mouse embryonic stem cells (mESCs) (Fraser et al., 2015) showed that dense L1 and B1 repeats appear to reside in distinct and mutually exclusive compartments across the mouse genome, whereas other types of retrotransposons such as ERV1, ERVK, and L2 tend to be randomly distributed (Figures 1A and S1B). The compartments marked by B1 repeats show enrichment of active histone marks (H3K4me3, H3K9ac, H3K27ac, H3K36me3), strong binding of RNA polymerase II (Pol II), and high levels of chromatin accessibility and transcription activity. In contrast, the compartments marked by L1 repeats show signatures of heterochromatin, including enrichment of the repressive H3K9me2 and H3K9me3 marks, and strong binding of heterochromatin proteins such as HP1α and the nuclear corepressor KRAB-associated protein-1 (KAP1 or TRIM28) (Figures 1A and S1B). Quantitative sequence analysis of annotated A/B compartments revealed high levels of SINE repeats (including B1, B2 and B4) in the A compartments and L1 repeats in the B compartments, whereas repeats in other subclasses showed no specific enrichment (Figures S1C, D). This opposing distribution pattern of B1/Alu and L1 in the A and B compartments was also detected in five other cell types (Figure 1B): human embryonic stem cells (hESCs), lymphoblastoid cells (GM12878) and erythroleukemic cells (K562), and mouse neural progenitor cells (NPCs) and neurons (Fraser et al., 2015; Rao et al., 2014). Furthermore, unsupervised clustering revealed that the genomic positions of A/B compartments in six distinct mouse and human cell types are highly similar to each other, with an average Spearman correlation coefficient >0.73 within species and >0.52 between species (Figure S1E). Compared to other subclass repeats, L1 and B1 are most strongly related to the high-order chromatin structures, and their distributions appear to be conserved in homologous regions of the mouse and human genomes (Figure S1E). For example, a region in mouse chromosome 2 (chr2: 120-140 Mb) and its syntenic region in human chromosome 20 (chr20: 6-50 Mb) show similar patterns of Hi-C contact probabilities, and gene and repeat compositions and distributions in the corresponding A and B compartments along the DNA sequences (Figure 1C). Thus, co-segregation of B1/Alu and L1 repeats with the A and B compartments appears to be invariable in different cell types in mouse and human.

**Figure 1.**
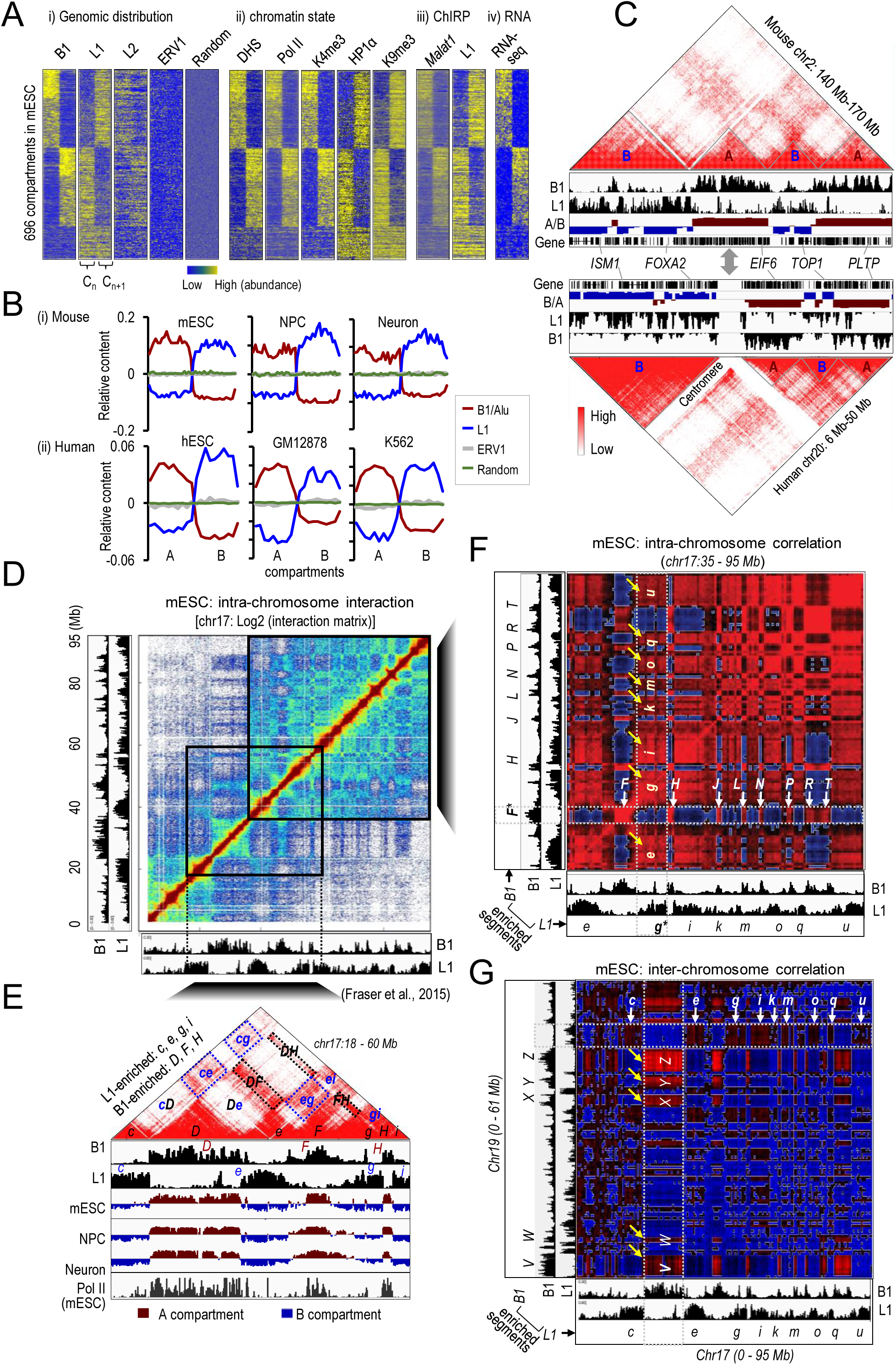
B1- and L1-rich genomic regions homotypically interact and correlate with A and B compartments. (A) Heatmaps of the distribution densities of B1, L1, L2, and ERV1 repeats and random genomic regions (panel i), DNase I hypersensitive sites (DHS) and ChIP-seq signals of Pol II, H3K4me3, HP1α, and H3K9me3 (panel ii), and ChIRP-seq signals of *Malat1* and L1 RNA (panel iii), and RNA-seq (panel iv) in mESCs across two adjacent compartments (Cn, Cn+1). All signals in 696 compartments annotated in mESCs were sorted according to the B1 distributions shown in panel (i). (B) Relative contents of various repeats across the A and B compartments annotated in various cell types in mouse (top) and human (bottom). Three repeat subclasses are shown (B1 or Alu, L1, and ERV1), together with random genomic regions as the negative control. (C) Genome browser shots showing conserved domain structures as indicated by heatmaps of the Hi-C contact matrix over a syntenic region in mouse (top) and human (bottom) ESCs. The B1 and L1 repeat densities, the A/B compartments shown by eigenvalues of the Hi-C contact matrix, and Refseq gene annotations are shown underneath each heatmap. (D) Heatmap of normalized interaction frequencies at 100-kb resolution on chromosome 17 in mESCs. Genomic densities of B1 and L1 repeats are shown at the left and bottom. (E) A zoomed-in view of the interaction matrix of the genomic region from 18 Mb to 60 Mb on mouse chr17 (40-kb resolution). Genomic densities of B1 and L1 repeats, eigenvalues of the Hi-C matrix representing A/B compartments from mouse mESCs, neural progenitor cells (NPC) and neurons, and Pol II ChIP-seq signals in mESCs are also shown underneath. B1-enriched regions are arbitrarily labeled as *D, F,* and *H* in uppercase. L1-enriched regions are labeled as *c, e, g,* and *i* in lowercase. Some strong homotypic interactions between compartments rich in the same repeat subfamily (for example, between the B1-rich regions *DF*, *DE* and *FH*, and between the L1-rich regions *ce* and *cg*), are highlighted by dotted boxes. (F) Correlation heatmap showing Pearson correlation coefficients of the interaction frequencies of any two paired regions in a sub-region on chr17 (500-kb resolution). B1-enriched regions are labeled in uppercase as *F, H … R, T*. L1-enriched regions are labeled in lowercase as *e, g … q, u*. Dotted boxes and arrows highlight positive correlations (in red) of the anchor region *F* (indicated by *\**) with other B1-enriched genomic regions (horizontal), and of the anchor region g (indicated by *) with other L1-enriched genomic regions (vertical). (G) Correlation heatmap showing Pearson correlation coefficients of inter-chromosomal interaction frequencies between chromosomes 17 and 19 at 500-kb resolution. B1-enriched regions on chr19 are arbitrarily labeled in uppercase as *V, W, X, Y,* and *Z*. L1-enriched regions on chr17 are labeled as shown in panels (E-F) from *c, e … q,* to *u*. The horizontal dotted box and arrows highlight positive correlations of an L1-enriched region on chr19 with other L1-enriched regions on chr17. The vertical dotted box and arrows show positive correlations of a B1-enriched region on chr17 with other B1-enriched regions on chr19.

We then took mouse chromosome 17 (chr17, 95 Mb in length) as an example to overlay L1 and B1 repeat features on the Hi-C interaction matrix of mESCs (Figures 1D-F). Interestingly, the plaid pattern of enriched and depleted interaction blocks in the Hi-C map is largely correlated with the compositions and distributions of B1 and L1 along the whole chr17 (Figure 1D). In a 42-Mb region of chr17, four L1-enriched B compartments (denoted by *c*, *e, g* and *i*) and three B1-enriched A compartments (denoted by *D*, *F* and *H*) are alternately positioned along the linear DNA sequence (Figure 1E). Strong interactions were observed between L1-enriched compartments (represented by *ce*, *cg, eg, ei* and *gi*) or B1-enriched compartments (represented by *DF*, *DH* and *FH*), but not between these two compartments (Figure 1E, dotted boxes). The interaction frequencies between *D* and *F* (*DF*) or between *c* and *e* (*ce*) are much stronger than those of *D* or *F* with *c* or *e* (*cD*, *De*, *eF*), despite the fact that these regions are closer in the linear sequence.

Conversion of Hi-C contact frequencies into Pearson correlation coefficients sharpened our view of the long-range chromatin interactions (Figure 1F). Strikingly, the plaid pattern of the Hi-C correlation map appears to perfectly match the distribution and interaction status of L1 and B1. L1-rich or B1-rich regions show strong enrichment of contacts with regions containing the same repeat type (red blocks in Figure 1F). We refer to these as homotypic contacts. Contacts between regions containing the other repeat type (heterotypic interactions) are strongly depleted (blue blocks in Figure 1F). For example, in one region of chr17 (35 to 95 Mb), L1-enriched segments (from *e* to *u*) and B1-enriched segments (from *F* to *T*) exhibit high frequencies of homotypic contacts (Figure 1F, highlighted by arrows), but strong depletion of heterotypic contacts. Similarly, homotypic contacts between L1-rich regions or B1-rich regions were also observed between chromosomes, as illustrated by chromosomes 17 and 19 (Figure 1G). These results indicate that genomic regions containing B1 or L1 repeats tend to interact with genomic regions containing repeat sequences from similar subfamilies, but not from different subfamilies, regardless of linear proximity. Thus, homotypic clustering of regions rich in B1 or L1 repeats partitions the genome into distinct A and B compartments, respectively.

### DNA FISH reveals the spatial segregation of L1 and B1 compartments

To visualize the nuclear localization of B1 and L1 repeats in cells we performed dual-color DNA fluorescence *in situ* hybridization (FISH) using fluorescence-tagged oligonucleotide probes that specifically target the consensus sequences of B1 and L1 elements. Consistent with the Hi-C analysis, L1 and B1 exhibit distinct and complementary nuclear localizations in mESCs (Figures 2A and 2B). While B1 DNA shows punctate signals in the nuclear interior, L1 DNA exhibits highly organized and concentrated signals that line the periphery of the nucleus and nucleolus. Weak L1 signals were also detected in a few areas of the nuclear interior subregions where B1 signals were absent. Both B1 and L1 signals are absent from DAPI-dense regions, which likely represent satellite repeat-enriched chromocenters (Mateos-Langerak et al., 2007). To confirm the L1 localization at the nucleolar periphery in mESCs, we performed immuno-FISH, which combines DNA FISH with immunofluorescence staining using an antibody against the nucleolar marker Nucleolin (NCL). Indeed, L1 signals surround and partially overlap with the ring-shaped signals of NCL at the nucleolar periphery (Figure 2C).

**Figure 2.**
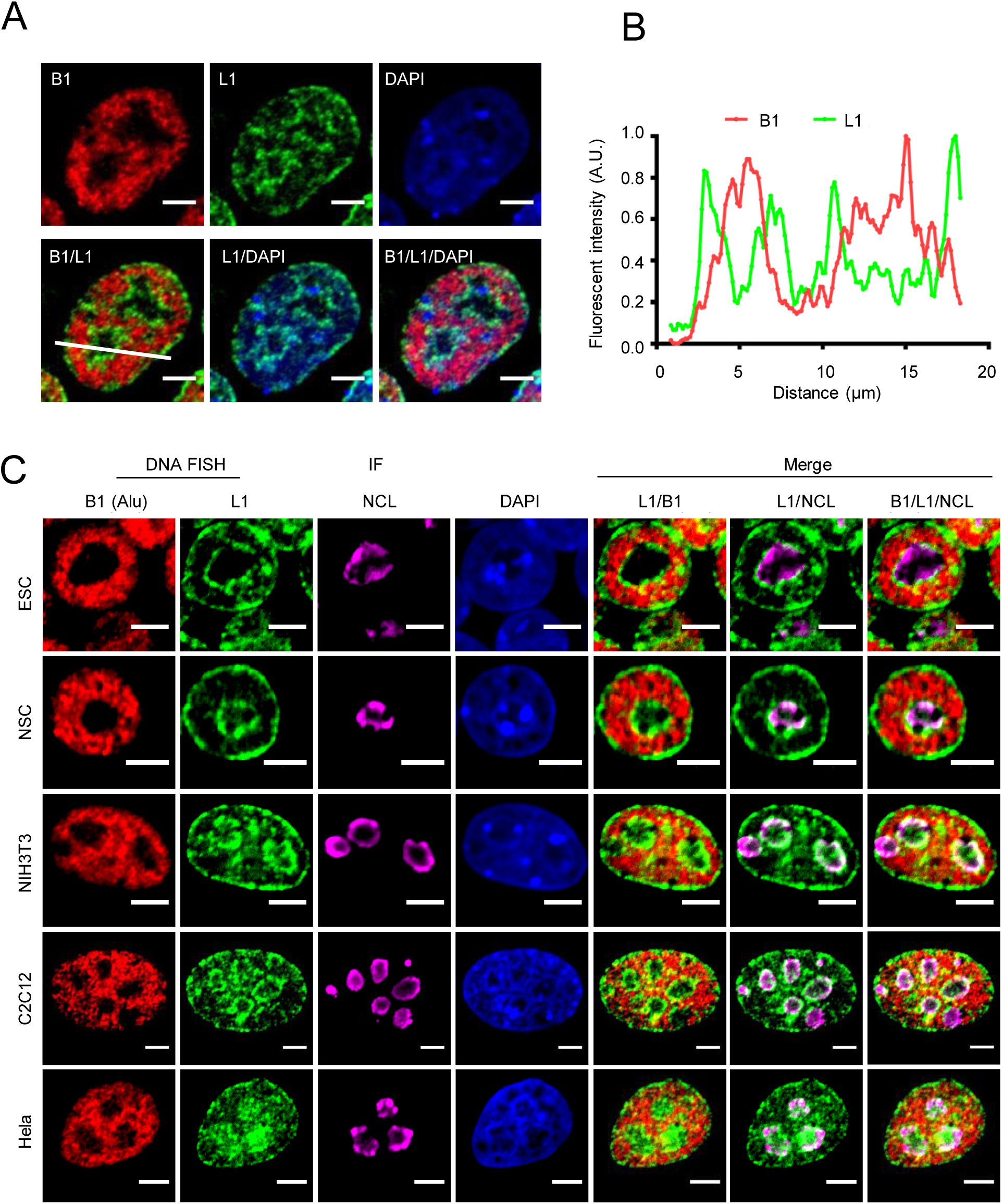
DNA FISH reveals the spatial segregation of L1 and B1 compartments. (A) Representative images of L1 (green) and B1 (red) repeats revealed by DNA FISH in mESCs. DNA is labeled by DAPI (blue). All scale bars, 5 μm. (B) Normalized fluorescence intensity profiles along the white line in the merged B1/L1 image shown in panel (A). The L1 channel shows four intensity peaks that surround the two intensity peaks in the B1 channel. These two peaks in the B1 channel correspond to valleys in the L1 channel. (C) Representative images of immuno-DNA FISH analysis of L1 (green) and B1 (red) DNA repeats, and NCL protein (purple) in mESCs (n=37), NSC (n=23), NIH3T3 (n=18), C2C12 (n=11) and Hela cells (n=15).

To ask whether L1 and B1 localizations might vary with cell type, we performed immuno-FISH in four other representative mouse and human cell lines: neural stem cells (NSC), fibroblasts (NIH3T3), myoblasts (C2C12) and Hela cells (Figure 2C). Similar to mESCs, all these cells show non-overlapping localizations of B1/Alu repeats, which are present in the nuclear interior, and L1 repeats, which are mainly located at the nuclear and nucleolar peripheries. The distributions of B1/Alu and L1 are reminiscent of the nuclear localization of euchromatin and heterochromatin, respectively. Together, the results from Hi-C and imaging analysis demonstrate that homotypic B1/Alu or L1 DNA sequences associate together to form larger clusters, which divide the nucleus into distinct territories of A/B compartments, and nuclear co-segregation of B1/Alu and L1 repeats with the A and B compartments is conserved across different cell types in mouse and human.

### Dynamic establishment of L1 and B1 segregation during the cell cycle and embryonic development

As chromatin organization undergoes dynamic changes during mitosis and after fertilization, we asked whether L1 and B1 compartments are established and re-constructed in these two biological processes. DNA FISH analysis of synchronous mESCs showed that L1 and B1 localizations change dramatically at different cell-cycle stages (Figures 3A and S2A). S-phase cells show non-overlapping and complementary localizations of L1 and B1 repeats (Figures 2 and 3A). This is similar to the pattern we observed previously in asynchronous mESCs, most of which are in the S phase of the cell cycle (Figure S2A). However, L1 and B1 DNA signals are mixed on mitotic chromosomes in metaphase (M phase, including prophase and anaphase), when the nuclear membrane and nucleoli are disassembled. As the cell cycle progresses into the G1 phase, L1 and B1 DNA start to segregate again. To quantify the degree of segregation, we defined a FISH-based segregation index as the negative value of Pearson’s correlation coefficient of L1 and B1 DNA signals in the nucleus. The FISH segregation index is lowest in M-phase cells, but increases significantly in the G1 phase and peaks in the S phase (Figure 3B).

**Figure 3.**
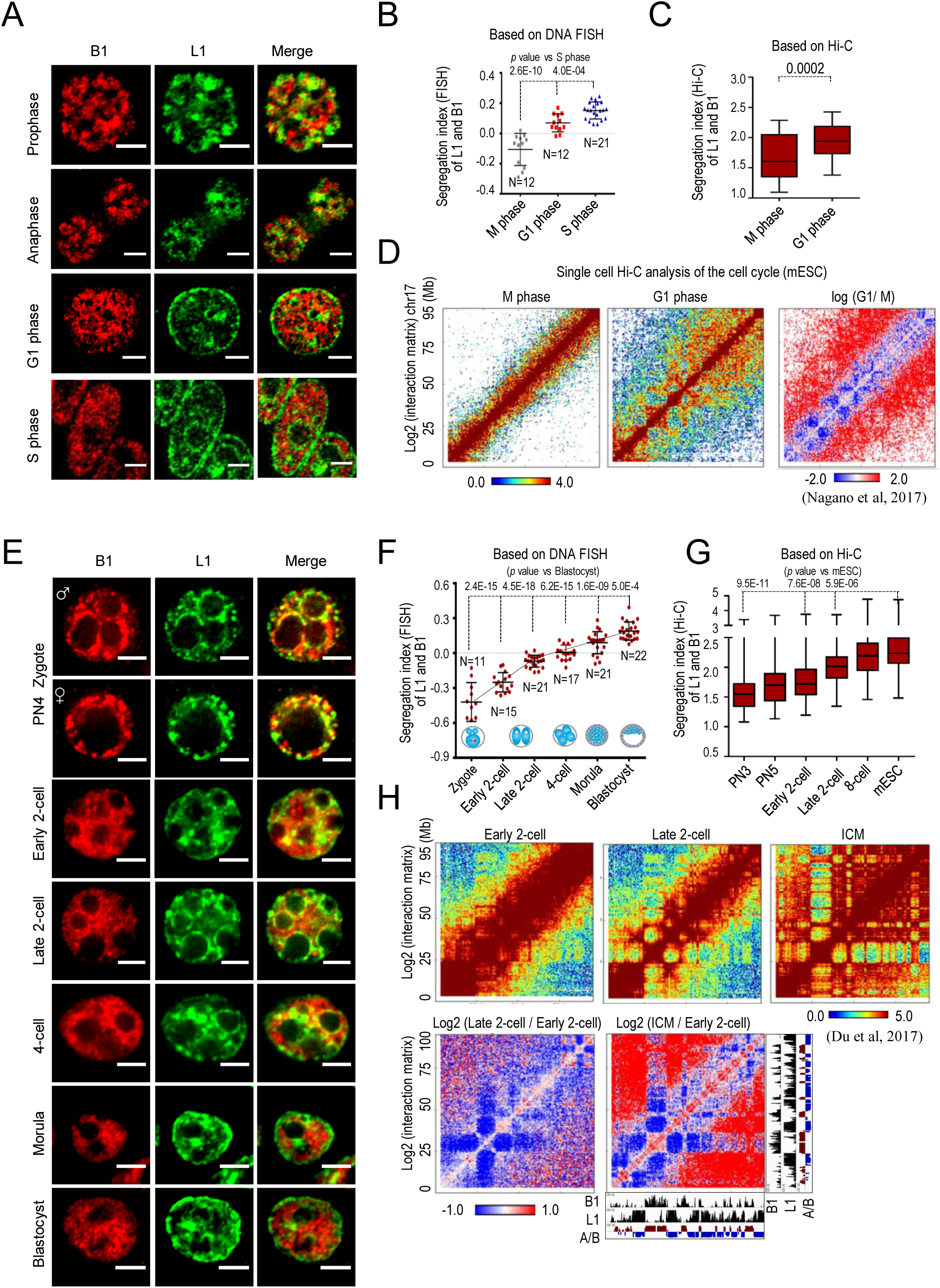
Dynamic segregation of L1 and B1 compartments during the cell cycle and embryonic development. (A) and (B) DNA FISH analysis of synchronized mESCs at different cell cycle stages. Representative images and the scatterplot of the segregation index of L1 and B1 DNA are shown in panels (A) and (B), respectively. Data are presented as mean ± standard deviation (SD). N, number of nuclei analyzed. M phase and G1 phase data are compared to S phase using the two-tailed Student’s *t*-test. *p* values are shown at the top. (C) Boxplot analysis of the ratio of homotypic interactions versus heterotypic interactions for B1 and L1 based on single-cell Hi-C data from mESCs. The *y*-axis shows the segregation index (Hi-C) of L1 and B1, which represents the ratio of average interaction frequency between compartments containing similar repeats (B1.B1 and L1.L1) to that between compartments containing different repeats (B1.L1) for all chromosomes (except X and Y). Larger values indicate a higher degree of homotypic interaction between B1- or L1-enriched compartments. *p* values are calculated with the two-tailed Student’s *t*-test. (D) Heatmaps of normalized interaction frequencies at 500-kb resolution on chromosome 17 in mESCs at M (left) and G1 (middle) phase. A comparison of contact frequencies between M and G1 phase [log2(G1/M)] for the whole of chromosome 17 is shown on the right. Genomic densities of B1 and L1 repeats are shown at the bottom. (E) and (F) DNA FISH analysis of early embryos. Representative FISH images and the scatterplot of the segregation index of L1 and B1 DNA are shown in panels (E) and (F), respectively. Data are presented as mean ± SD (embryos were collected and processed in two independent experiments). N, number of nuclei analyzed. Each sample is compared to the blastocyst stage and *p* values are calculated with the two-tailed Student’s *t*-test. Dotted lines show the trend-line. (G) Boxplot showing a gradual increase of homotypic versus heterotypic interactions between L1-rich and B1-rich regions based on Hi-C analysis of early embryos from zygotes (including PN3 and PN5 stages) to 8-cell embryos and pluripotent mESCs (*in vitro* equivalent of the inner cell mass cells of blastocysts). Each sample is compared to mESCs and *p* values are calculated with the two-tailed Student’s *t*-test. (H) The upper panels show heatmaps of normalized interaction frequencies at 500-kb resolution on chromosome 17 in mouse embryos at the early 2-cell, late 2-cell and inner cell mass (ICM) stages. The lower panels show comparisons of the contact frequencies between Late 2-cell and Early 2-cell (left), and ICM and Early 2-cell (right). Genomic densities of B1 and L1 repeats are also shown.

To provide further molecular evidence for segregation of repeats during the cell cycle, we analyzed the published Hi-C data from cell-cycle synchronized mESCs and HeLa cells (Nagano et al., 2017; Naumova et al., 2013). In both cell types, G1-phase cells exhibit a classic plaid pattern of hierarchical interactions, with enriched and depleted interaction blocks outside of the diagonal region of the Hi-C interaction heatmap (Figures 3C, 3D, S2B, and S2C). In contrast, M-phase cells exhibit stronger signals along the diagonal, which represent the linearly organized, longitudinally compressed array of consecutive chromatin loops. To quantify this difference, we defined a Hi-C-based segregation index by calculating the ratio of homotypic versus heterotypic interaction frequencies between L1 and B1/Alu subfamilies. Indeed, the Hi-C segregation index is significantly higher in G1-phase cells than in M-phase cells (Figures 3C and S2B). These results indicate that segregation of B1/Alu and L1 repeats is dispersed by mitosis, and is re-established when the higher-order chromatin structure forms during each cell cycle. This finding agrees with previous reports that in metaphase, chromosome folding becomes homogeneous and large megabase-scale A and B compartments are lost, whilst in interphase, chromosomes return to a highly compartmentalized state (Nagano et al., 2017; Naumova et al., 2013).

After fertilization, the chromatin undergoes extensive reprogramming from a markedly relaxed state in zygotes to fully organized structures in blastocysts (Du et al., 2017; Flyamer et al., 2017; Ke et al., 2017). We performed time-course DNA FISH analysis of L1 and B1 in early mouse embryos (Figure 3E). L1 and B1 signals are largely overlapping in zygotes and early 2-cell embryos, and become progressively more segregated during embryonic divisions (Figure 3E). A plot of the FISH segregation indexes along the course of blastocyst development showed that the greatest change (steepest trend-line) occurred between the zygote and the late 2-cell stage (Figure 3F). This observation implies that the initiation of B1 and L1 compartmentalization may coincide with the zygotic genome activation (ZGA), during which massive transcription switches on (Qiu et al., 2003). We then analyzed the published Hi-C data from early mouse embryos (Du et al., 2017). Consistent with our results, we detected a gradual increase of the segregation index (Figure 3G). Early 2-cell embryos exhibit prevalent *cis*-chromosomal contacts along the diagonal of the Hi-C interaction map, whereas the plaid patterns of Hi-C interactions become readily detectable in late 2-cell embryos and are fully established in the inner cell mass (ICM) cells of blastocysts (Figure 3H). Thus, in early embryos, compartmentalization of B1- and L1-rich regions appears to be established in a stepwise manner, coincident with *de novo* establishment of higher-order chromatin structures.

### Repeat RNA and transcription promote L1 and B1 compartmentalization

Transcription and nuclear architecture are closely interdependent and influence each other (Dowen et al., 2014; Rowley and Corces, 2018; Rowley et al., 2017). We sought to test whether RNA transcribed from L1 and B1 repeats mediates the function of their DNA sequences. We first analyzed the effects of inhibition of L1 and B1 expression in early mouse embryos, as both repeats are activated and highly expressed in two-cell embryos (Fadloun et al., 2013; Jachowicz et al., 2017; Rothstein et al., 1992). We recently showed that that L1 RNA can be efficiently knocked down with antisense morpholinos (AMO) in mouse embryos and mESCs (Percharde et al., 2018). We successfully depleted L1 and B1 transcripts by injecting antisense morpholinos or oligonucleotides (ASO) into mouse zygotes (Figures S3A-B). Embryos depleted of L1 or B1 RNA were able to pass the first embryonic division and their development was indistinguishable from control embryos at the 2-cell stage; however, they failed to divide further and became arrested at the 2-cell stage (Figure 4A). We collected these embryos for DNA FISH analysis when the control group injected with scramble AMO or ASO had grown to the late 2-cell stage. In embryos depleted of L1 or B1 RNA, L1/B1 segregation is impaired, as indicated by significantly lower L1/B1 segregation indexes compared to scramble control and noninjected late 2-cell embryos (Figures 4B and S3C-E). This result suggests that formation of L1 and B1 compartments is delayed upon depletion of L1 or B1 transcripts, which is consistent with the observed embryonic arrest.

**Figure 4.**
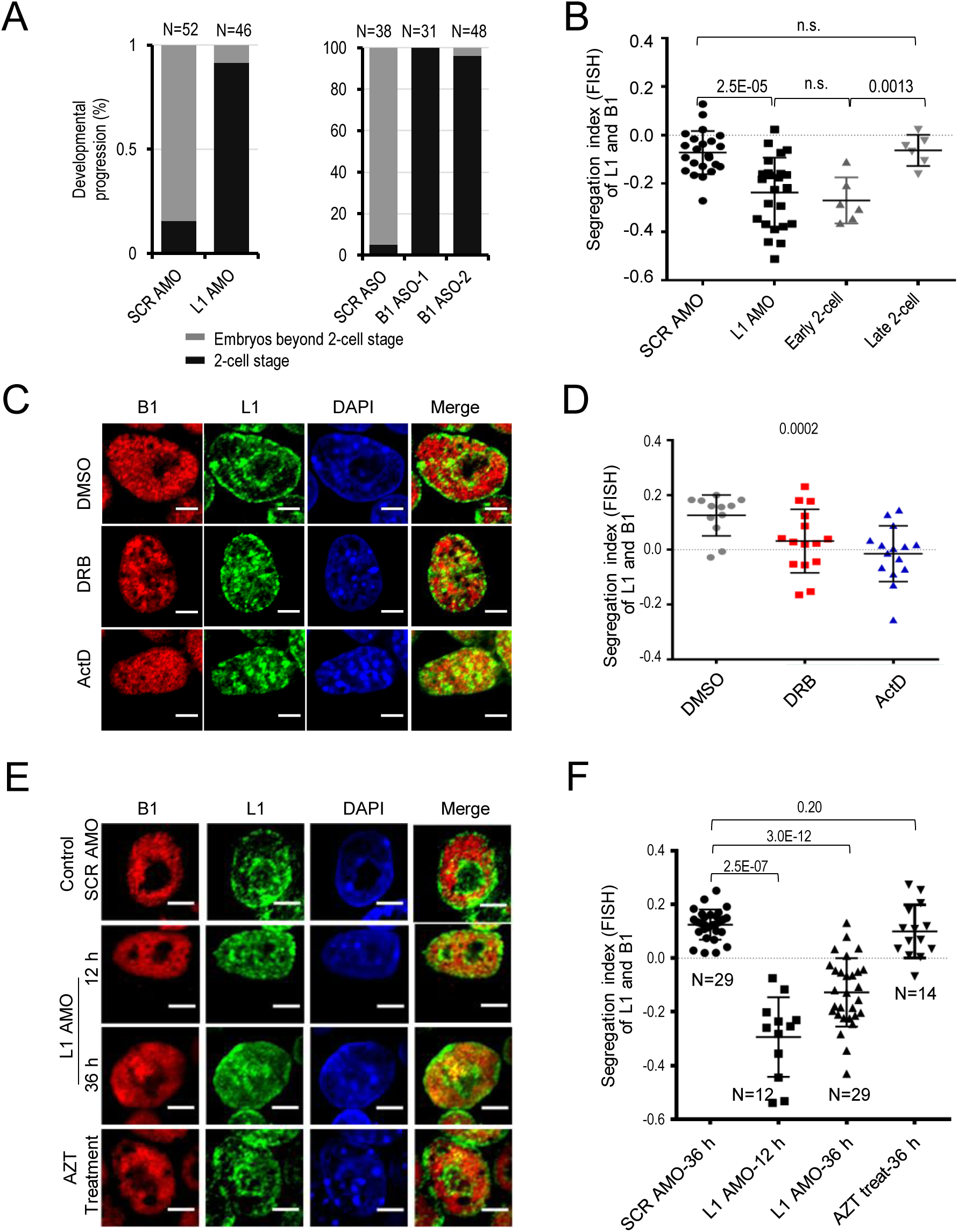
Repeat RNA and transcription promote the spatial segregation of L1 and B1 compartments. (A) Developmental analysis of embryos microinjected with scramble or L1 AMO (left) or with scramble or two different B1 ASOs (ASO-1 and ASO-2) (right) at the zygote stage. N, number of embryos analyzed. (B) Embryos depleted of L1 RNA by AMO show poorer L1/B1 segregation, as indicated by significantly lower L1/B1 segregation indexes compared to the scramble AMO control and noninjected late 2-cell embryos. Each dot represents an embryo analyzed. Black and grey dots represent injected and noninjected embryos, respectively. Data were collected from three independent experiments. *p* values are calculated with the two-tailed Student’s *t*-test. (C) and (D) DNA FISH analysis of L1 (green) and B1 (red) repeats in mESCs treated with transcription inhibitors for 3 hours. Representative images and the scatterplot of the segregation index of L1 and B1 DNA are shown in (C) and (D), respectively. DMSO: mock control; DRB (100 μM): inhibits the release and elongation of Pol II; ActD (1 μg/ml): inhibits both Pol I and II. Both drug treatments disrupt the perinucleolar staining of L1 DNA and induce mixing of the L1 and B1 DNA signals. Treatment with ActD elicits a stronger mixing effect than DRB. (E) and (F) DNA FISH analysis of L1 (green) and B1 (red) repeats in mESCs transfected with scramble AMO for 36 hours, or L1 AMO for 12 hours and 36 hours, or treated with the drug AZT for 12 hours. Representative images and the scatterplot of the segregation index of L1 and B1 DNA are shown in (E) and (F), respectively. All scale bars, 5 μm. N, number of nuclei analyzed. Data are presented as mean ± SD (two independent experiments), and *p* values are calculated with two-tailed Student’s *t*-test.

It was reported that inhibition of Pol II transcription by α-amanitin causes embryonic arrest at the late 2-cell stage (Du et al., 2017; Qiu et al., 2003). We analyzed the Hi-C data obtained by Du *et al* and found that α-amanitin-treated embryos exhibit predominantly *cis*-chromosomal contacts along the diagonal of the Hi-C map, in a pattern similar to that of early 2-cell embryos, while the control group shows the plaid-like pattern of late 2-cell embryos (Figures S4A). Consistent with this, treating mESCs with the transcription inhibitors Actinomycin D (ActD) or DRB (5,6-dichloro-1-b-D-ribofuranosylbenzimidizole) also altered L1/B1 compartmentalization as shown by loss of L1 perinucleolar localization and gain of mixed L1 and B1 signals (Figures 4C, 4D and S4B). Notably, inhibition of both Pol I and II by ActD caused a more severe effect than inhibition of Pol II alone by DRB. These results indicate that transcription facilitates the nuclear organization.

Next, we sought to deplete repeat transcripts in mESCs in order to dissect the effects on chromatin organization independent of embryonic progression. B1/Alu repeats are known to be broadly involved in diverse cellular processes, including transcription and RNA processing and nuclear export (Chen et al., 2009; Dominissini et al., 2011; Lubelsky and Ulitsky, 2018; Polak and Domany, 2006). Treatment of mESCs with B1 ASO led to severe cell death within hours of transfection (data not shown), which precluded direct assessment of the function of B1 transcripts in organizing the nuclear structure. L1 repeats, the most abundant of all repeat subclasses, are predominantly enriched in heterochromatic B compartments. It was suggested that heterochromatin interactions play a more important role in nuclear compartmentalization than euchromatin interactions (Falk et al., 2019; Houda Belaghzal, 2019). To reveal a direct role of repeat RNA in the nuclear organization, we focused on L1 in subsequent investigations.

Treatment of mESCs with L1 AMOs drastically altered the formation of L1/B1 compartments as shown by the appearance of the overlapping, yellowish FISH signals of B1 and L1 DNA(Figures 4E, 4F, and S5A). In particular, when L1 RNA was depleted by AMO treatment for 12 or 36 hours, perinucleolar L1 signals were diffuse or absent and nucleoplasmic signals were dramatically increased at both time-points. In contrast to the punctate pattern of B1 DNA in the control mESCs, B1 signals became more uniformly dispersed in the nucleoplasm, which is indicative of decreased clustering. In comparison, treatment with the drug azidothymidine (AZT), which blocks L1 retrotransposition activity (Xie et al., 2010), failed to affect L1 RNA levels and the nuclear localization of L1 and B1 (Figures 4E, 4F, and S5B). In addition, statistical analysis indicated that the FISH segregation indexes were significantly lower in L1-depleted mESCs than in mESCs treated with scramble AMO or AZT. These results suggest a role of L1 transcripts in regulating the nuclear localization and segregation of L1 and B1 DNA repeats, independently of L1 retrotransposition activity, in mESCs.

### L1 RNA regulates the higher-order chromatin structure in mESCs

For in-depth characterization of the molecular defects caused by L1 RNA depletion, we performed *in situ* Hi-C analysis at 36 hours after transfecting L1 AMO into mESCs. Direct visualization of Hi-C interaction maps revealed obvious differences in the plaid pattern of L1-depleted and control mESCs, as illustrated by chr17 (Figure 5A). In L1-depleted mESCs, Hi-C contact signals were abnormally increased along the diagonal line, whereas the plaid signals outside of the diagonal regions became fuzzy or even lost (Figure 5A, panel i). Accordingly, comparison of Hi-C contact frequencies of the control versus L1-depleted cells showed decreased ratios (in blue) across the diagonal and increased ratios (in red) in the periphery (Figure 5A, panel ii). These changes indicate enhanced local chromosomal contacts but decreased long-range interactions in L1-depleted mESCs.

**Figure 5.**
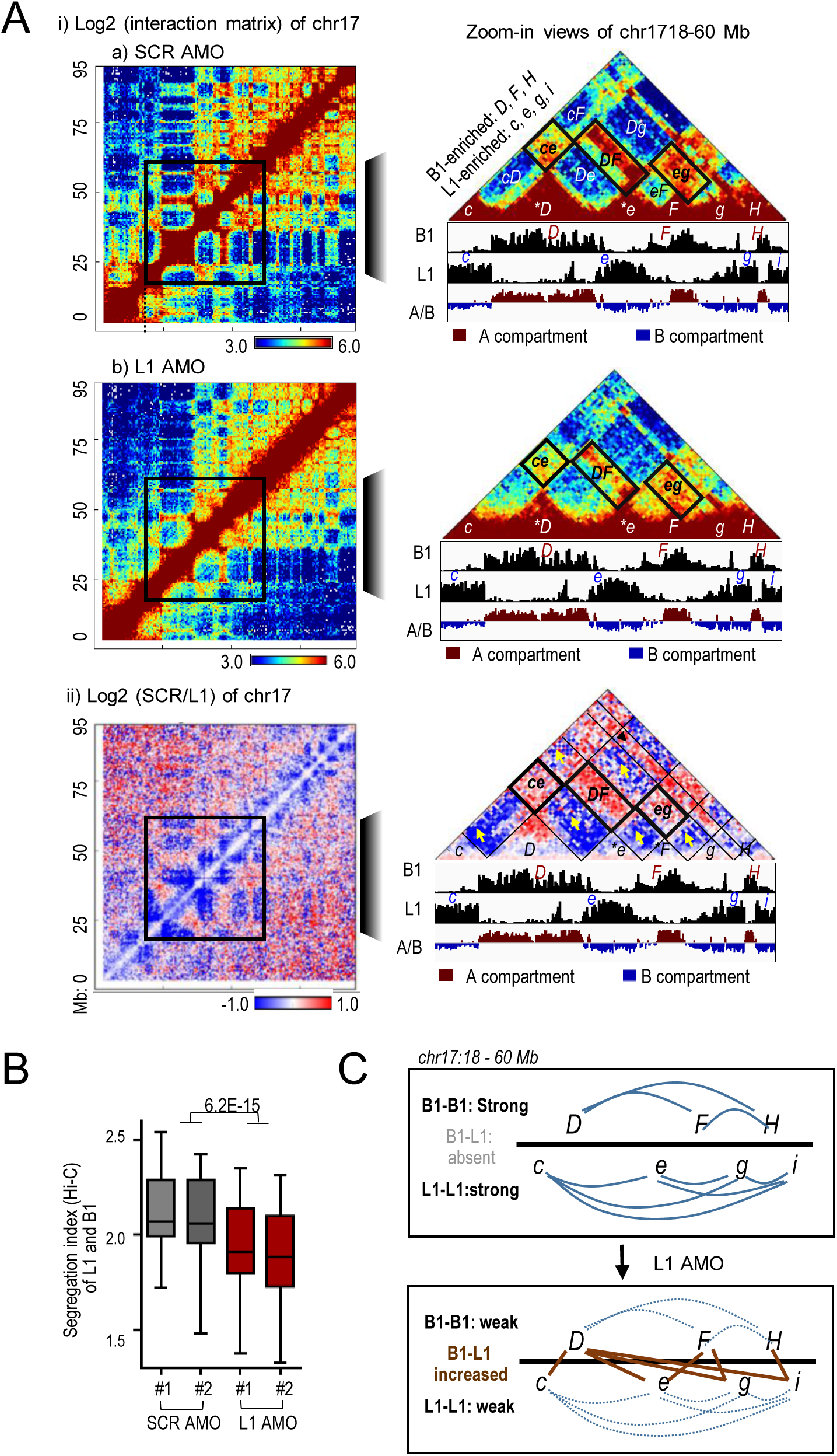
L1 RNA is required for the formation and maintenance of higher-order chromatin structure. (A-B) Hi-C analysis of mESCs treated with scramble control (SCR) and L1 AMOs for 36 hours. Two independent experiments of AMO transfection followed by Hi-C were performed. Panel A shows Hi-C heatmaps of contact frequencies for SCR and L1 AMO (panel i) and the comparison of contact frequencies (panel ii) between scramble and L1 AMOs [log2(SCR/L1)]. The whole of chromosome 17 (chr17) is shown on the left (at 500-kb resolution) and the boxed sub-region (18-60 Mb) of chr17 is shown enlarged on the right. To better orient the visualization and comparison of these three sub-region heatmaps, representative homotypic interactions (*ce*, *DF*, and *eg*) are labeled with black boxes in the right panels. Relevant B1- and L1-enriched compartments are labeled as in Figure 1. Panel B shows a boxplot comparing the Hi-C-based segregation indexes. L1 AMO led to decreased ratios of homotypic interaction versus heterotypic interaction between L1-rich and B1-rich regions compared to SCR AMO. The *p* value was calculated by the two-tailed Student’s *t*-test. (C) Schematic representation of compartmental interactions before and after depletion of L1 RNA based on Hi-C data shown in the panel (A). The control mESCs show strong homotypic interactions (indicated by blue solid lines), whereas L1-depleted cells show weakened homotypic interactions (dotted blue lines) and abnormal increases of heterotypic interactions (indicated by brown solid lines).

A zoomed-in view of a 42-Mb region of chr17 in L1-depleted mESCs further shows decreased homotypic chromatin contacts between L1-rich or B1-rich compartments, but abnormally increased heterotypic contacts (Figure 5B, right; and Figure 5C). For example, B1-B1 interactions (represented by *DF*, *DH* and *FH)* or L1-L1 interactions (represented by *ce*, *cg*, *ci, eg, ei* and *gi)* were downregulated in L1-depleted cells, whereas aberrant B1-L1 contacts (represented by *cD*, *De*, *Dg, Di, eF, Fg,* and *Hi*) were increased. Globally, L1-depleted cells show significantly lower Hi-C segregation indexes compared to control mESCs (Figure 5A). Consistent with the overlapping signals of L1 and B1 DNA FISH (Figure 4E), these results revealed defective nuclear segregation of L1 and B1 compartments after depletion of L1 RNA (Figure 5B).

Next, to directly visualize changes in spatial contacts at specific loci, we performed oligopaint dual-color DNA FISH (Figure 6A). We labeled four representative genomic regions on chr17, a B1-enriched region (*F*) and three L1-enriched regions (*e*, *g* and *q*), with a set of 500 ∼ 4,500 DNA probes for each region at a density of 200 bp per probe, and measured the spatial distances between the probe signals within mESC nuclei. In control mESCs, the two linearly adjacent L1- and B1-enriched regions *e* and *F* are positioned away from each other in the nuclear space with a median distance of 1.72 ± 0.55 µm. Depletion of L1 RNA significantly shortened the distance between them to 1.27 ± 0.47 µm (Figures 6B and 6C, left). In comparison, the two L1-enriched compartments *g* and *q* reside in close spatial proximity (1.08 ± 0.45 µm) in the nuclei of control cells even though they are separated by multiple L1 and B1 compartments in the linear genomic sequence. L1 depletion significantly increased the nuclear distance between *g* and *q* to 1.81 ± 0.82 µm (Figure 6C, right). Together, DNA FISH and Hi-C results demonstrate that L1 RNA is critical in maintaining the high-order chromatin structure of segregated L1- and B1-enriched compartments in mESCs.

**Figure 6.**
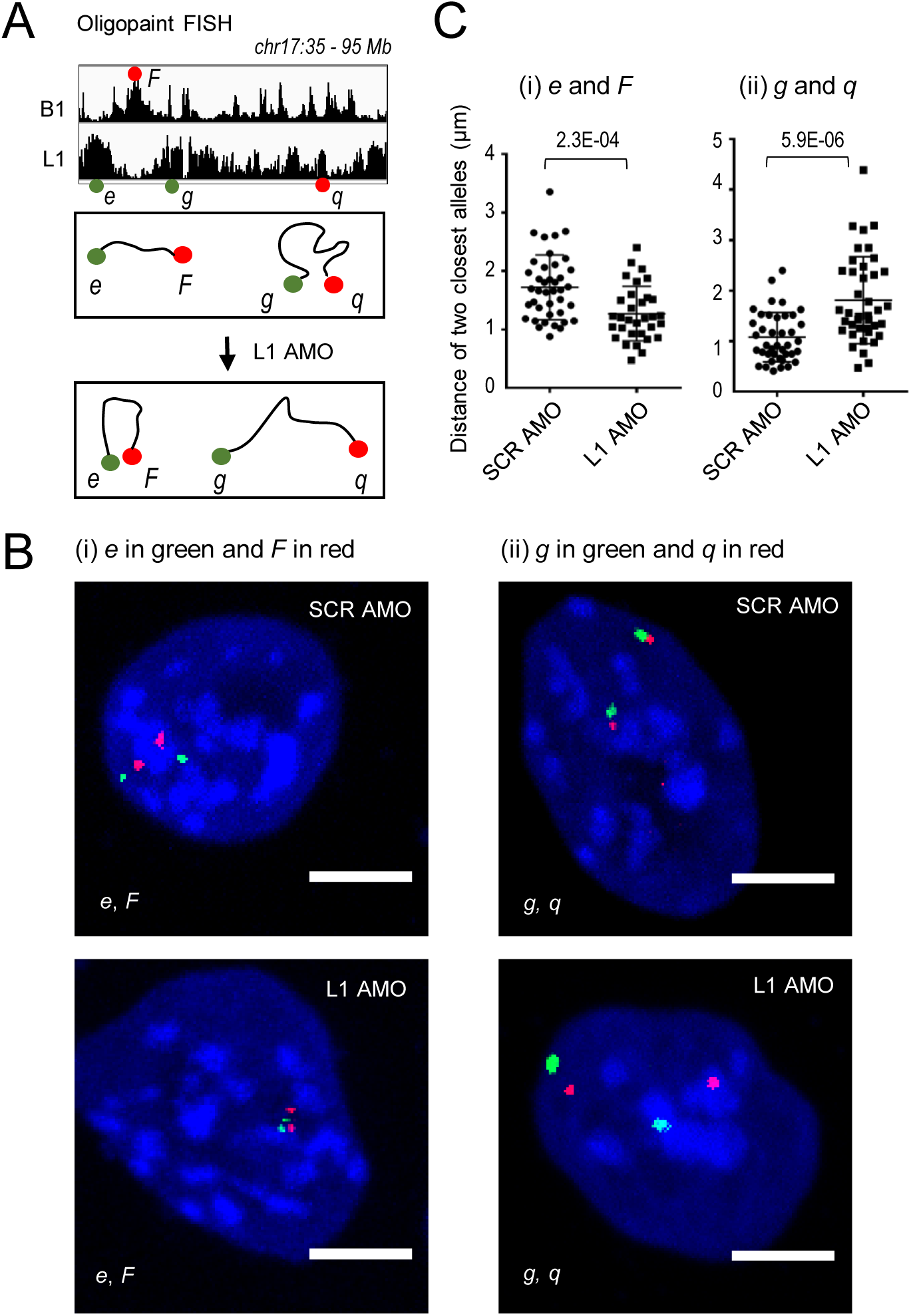
L1 RNA is required for spatial segregation of L1 and B1 compartments. Oligopaint DNA FISH analysis of four representative loci before and after depletion of L1 RNA for 36 hours in mESCs. (A) Scheme of relative chromosomal localizations and spatial distances of the four compartments labeled by Oligopaint DNA FISH probes. B1-enriched domain: *F*. L1-enriched domains*: e, g,* and *q.* Representative images and quantification of the spatial distances between two labeled compartments are shown in panels (B) and (C), respectively. Each dot in panel (C) represents a nucleus. In cells treated with L1 AMO, the median distance between the two B1 and L1 compartments *e* and *F* is significantly decreased compared to the control, while the median distance between the two L1 compartments *g* and *q* is significantly increased. Data are presented as mean ± SD. N, number of nuclei analyzed. *p* values are calculated by the two-tailed Student’s *t*-test. Scale bars, 5 μm.

### L1 repeats promote phase separation of HP1α

Recent evidence suggests that the formation of heterochromatin domains entails a phase-separation mechanism (Larson et al., 2017; Li et al., 2012; Strom et al., 2017; Zwicker et al., 2014). HP1α, a known component of heterochromatin, forms phase-separated droplets in the presence of DNA or nucleosomes *in vitro* (Larson et al., 2017; Strom et al., 2017). We showed that HP1α binds strongly to L1-enriched but not B1-enriched genomic regions (Figure 7A). In addition, L1 RNA specifically binds L1 DNA sequences throughout the genome, and is significantly enriched in L1 repeat-associated compartments, but is depleted in B1-associated compartments (Figures 1A and 7A), as shown by chromatin isolation by RNA purification followed by sequencing of L1 RNA (ChIRP-seq) (Lu et al., paper submitted). Given the abundance and co-residence of L1 repeats with HP1α in similar heterochromatin contexts within the nucleus, we hypothesized that L1 RNA may facilitate heterochromatin formation by promoting HP1α phase separation.

**Figure 7.**
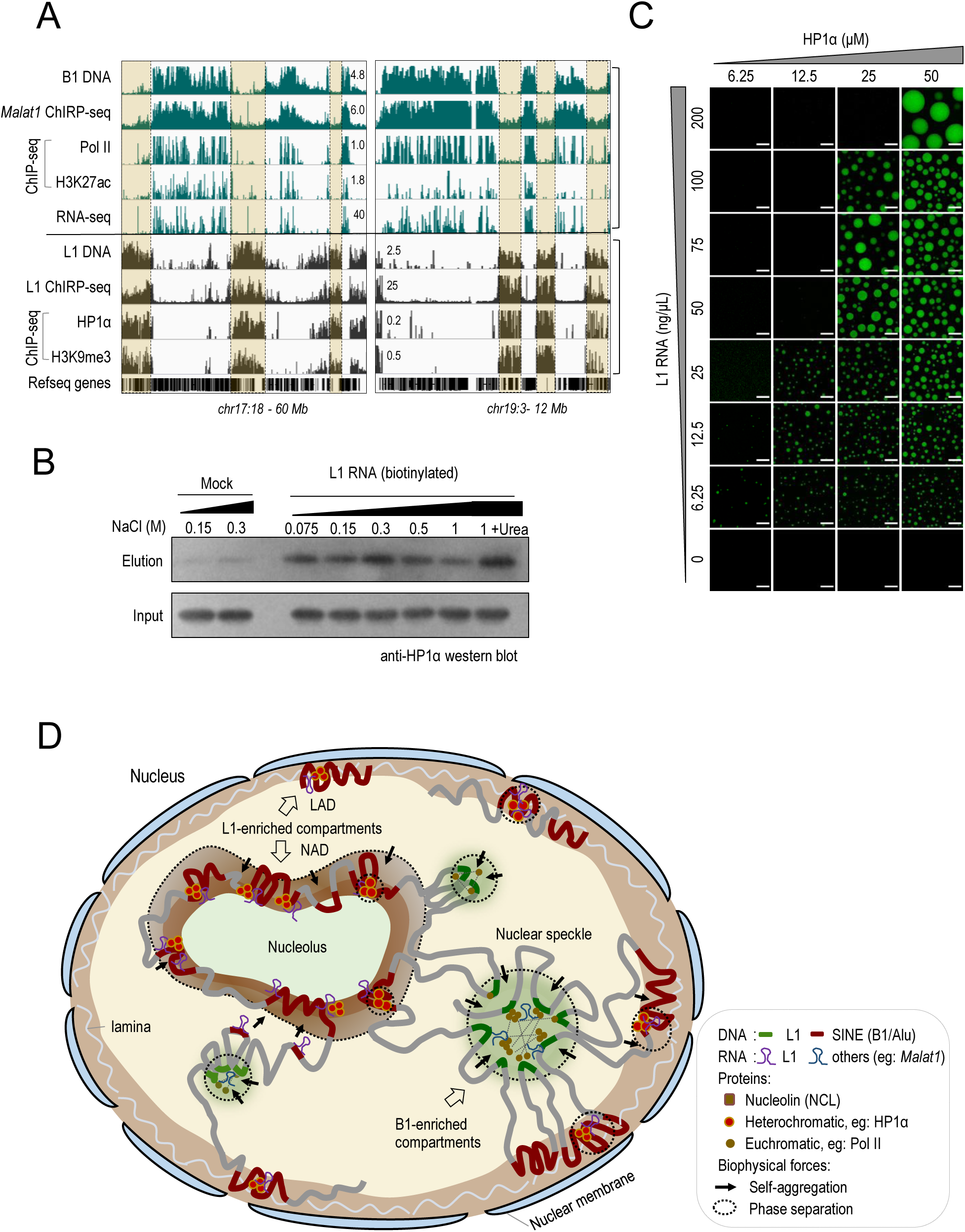
L1 RNA binds to its own DNA and promotes the phase separation of HP1α. (A) Genome browser view of mouse chr17: 18 - 60 Mb (left) and chr19: 3 - 12 Mb (right). The first five rows show the genomic density of B1 repeats, B1-related sequencing data (ChIRP-seq signals of *Malat1,* ChIP-seq signals of Pol II, and H3K27ac), and RNA-seq in mESCs. The lower tracks show the genomic density of L1 repeats (highlighted by beige shading) and L1-related sequencing data, including ChIRP-seq signals of L1 RNA, ChIP-seq signals of HP1α and H3K9me3. Refseq gene annotations are also included. (B) Biotinylated L1 RNA pulls down recombinant HP1α protein. Reactions without addition of L1 RNA were served as mock control. (C) Representative images of droplet formation at different concentrations of HP1α protein and L1 RNA. Concentrations of HP1α and RNA are indicated at the top and left of the images, respectively. Scale bars, 10 μm. Data are representative of 3 independent experiments. (D) The model. First, the intrinsic self-assembly property of L1 and B1/Alu repeats provides numerous nucleation points to seed the formation of nuclear subdomains. Repetitive DNA sequences also serve as anchor sites for transcription machinery, regulatory proteins and RNAs. Second, the embedded structural information in DNA repeats may be translated by their RNA transcripts, together with interacting DNA and/or RNA-binding proteins. Molecular crowding generated by interactions among DNA sequences, RNAs, and proteins in subnuclear domains seeded by individual clusters of L1 or B1/Alu, may drive the aggregation of compartments containing the same repeat type through a phase-separation mechanism, which consequently folds the genome. Third, the nuclear segregation of L1-enriched B compartments and B1/Alu-enriched A compartments may be further reinforced by attaching their DNA sequences to subnuclear structures such as nuclear speckles and the nucleolus, respectively, which serve as scaffolds to stabilize the nuclear architecture.

We firstly tested the direct protein-RNA interaction between the recombinant human HP1α protein and L1 RNA fragments that were transcribed *in vitro*. Interestingly, the L1 RNA mix captures HP1α protein in extremely high stringent washes (up to 1 M of salt and urea), indicating that HP1α has a strong RNA-binding activity towards L1 (Figure 7B). Droplet formation assays showed that HP1α alone fails to phase separate (Figures 7C), consistent with a previous report (Larson et al., 2017). Intriguingly, upon addition of L1 RNA, HP1α forms spherical droplets with the properties of liquid-like condensates, such as fusion of droplets and rapid recovery after photo-bleaching (Figures 7C and S6A, B). HP1α-L1 RNA droplets increase in size when increasing concentrations of components are added into the assay (Figure 7C). L1 DNA also promotes the phase separation of HP1α (Figure S6C). These results suggest that L1 repeats play an active role in heterochromatin formation.

## Discussion

Although tremendous efforts have been dedicated to studies of structural chromatin proteins and cataloging chromatin maps, the role of DNA sequences in 3D genome organization has been largely ignored. It has been speculated that genome organization may occur through polymer phase separation mediated by homologous sequence pairing (Cook, 2002; Cook and Marenduzzo, 2018; Falk et al., 2019; Rowley et al., 2017; Tang, 2017). However, it was unclear which types of repeats are involved in this process, and direct evidence was still lacking. In addition, the molecular determinants and mechanisms that drive formation of the A/B compartments remain unclear. In this study, we provide experimental evidence to support a fundamental role of L1 and B1/Alu repeats in organizing chromatin macrostructure.

Complementary results based on genomics, Hi-C and imaging analysis revealed that L1 and B1/Alu repeats tend to cluster with sequences from their own repeat subfamily and form mutually exclusive domains in the nuclear space that are highly correlated with A/B compartments (Figures 1 and 2). The nuclear segregation of B1/Alu-enriched A compartments in the nuclear interior and L1-enriched B compartments in the nuclear and nucleolar peripheries is relatively conserved and stable across a variety of mouse and human cells, and re-occurs during the cell cycle (Figure 2). In addition, *de novo* establishment of the segregated pattern of L1- and B1-rich compartments is coincident with the formation of higher-order chromatin structures during early embryogenesis, and appears to be critically regulated by RNA transcripts produced from L1 and B1 DNAsequences. Depletion of L1 RNAby AMOs in mESCs drastically disrupts the nuclear localization of L1 repeats, and leads to impaired segregation of L1 and B1 compartments and altered higher-order chromatin structures. Moreover, L1 RNA binds to its own DNA sequences in cells, and promotes HP1α phase separation *in vitro*. Together, our findings suggest that repeat transcripts may transact the structural information embedded in their DNA sequences and facilitate formation of the nuclear architecture with assistance from protein partners, such as HP1α (Figures 4-7).

Based on our observations together with previous reports in the literature, we propose a hypothetical model in which repetitive elements organize the macroscopic structure of the genome at three hierarchical levels (Figure 7D). First, L1 and B1/Alu repeats serve as the genetic basis for A and B compartments. The abundance and scattering of these repeat elements in the genome provide numerous nucleation points or “structural codes” to seed the formation of nuclear subdomains. Homotypic clustering of Ll-rich regions or B1-rich regions initiates genome folding. Second, the structural information embedded in linear genomic DNA repeats is in part transacted by their transcripts, particularly L1 RNA as demonstrated in this study, into spatially ordered chromatin in the nucleus. Molecular crowding and specific interactions among similar molecules, including DNA sequences and RNAs containing multiple copies of the same repeat type, and their associated proteins, promote clustering into larger subdomains (Bergeron-Sandoval et al., 2016; Cook, 2002; Misteli, 2007). Phase separation of individual subdomains may also lead to their segregation in the nuclear space and the formation of distinct L1-enriched heterochromatin and B1/Alu-enriched euchromatin domains. Third, chromatin compartmentalization may be further reinforced and stabilized through attachment of repeat DNA sequences to subnuclear structures such as the nucleolus and nuclear speckles (Quinodoz et al., 2018). Evidence to support this notion includes our observation that L1 repeats are preferentially localized at the nuclear and nucleolar peripheries, and depletion of L1 RNA disrupts the localization of L1 DNA at these sites.

In the absence of zygotic transcription, chromatin folding continues to occur in a significantly delayed manner (Figures 4 and S4), and the higher-order chromatin structure still partially forms (Du et al., 2017). This observation implies that the 3D organization of chromatin is an autonomous process that is greatly facilitated by transcription. We surmise that homotypic clustering of L1-rich or B1-rich regions is autonomous but dynamically regulated. L1/B1-driven nuclear compartmentalization is reminiscent of the self-assembly of tandem repeat sequences such as ribosomal DNA (rDNA) and satellite repeats, which promotes high-order assemblages of the nucleolus and pericentromeric domains, respectively (Falahati et al., 2016; Tjong et al., 2016). Consistent with our model, in mouse ESCs carrying a human artificial chromosome, human L1 sequences spatially interact with mouse L1-enriched genomic regions but avoid the SINE-enriched regions, and *vice versa* for human SINEs (van de Werken et al., 2017). Specific DNA-RNA-protein interactions at homologous L1 or B1 chromosomal segments not only enhance the formation of subdomains through phase separation, but may also promote their segregation in the nucleus. A recent study reported that there does not appear to be cross-talk between different transcription factors; instead, they interact among themselves via sequence-specific protein-protein interactions to form concentrated hubs on synthetic lacO DNA arrays (Chong et al., 2018). L1 or B1/Alu repeats tend to associate with distinct sets of transcription and chromatin regulators (Lu et al., paper submitted). For example, heterochromatin proteins such as HP1α, KAP1, and SETDB1 are specifically enriched on L1 elements, whereas general transcription factors such as GTF3C2 and CEBPB and RNA Pol II subunits show enriched binding on B1/Alu. In addition, different physiochemical properties of phase-separated L1 or B1 subdomains, for example chromatin compactness, may further promote the segregation of L1 and B1 compartments.

By focusing on L1 transcripts, we demonstrate an essential role for L1 repeat RNA in promoting the formation of segregated higher-order chromatin structure in mESCs (Figures 4-6). Heterochromatin domains appear to be more stable than euchromatin domains (Houda Belaghzal, 2019). In support of a heterochromatin-dominated model (Falk et al., 2019), our data further suggest that L1-mediated heterochromatin formation may drive nuclear compartmentalization. This function of L1 RNA might be achieved by stabilizing the binding of chromatin proteins such as HP1α to L1 DNA sequences, and by recruiting additional RNA-binding proteins to increase local protein concentrations and multivalent interactions for efficient phase separation. It was reported that the RNA-binding activity of mammalian HP1α is required for its localization to pericentromeric chromatin (Muchardt et al., 2002; Stunnenberg et al., 2015). On the other hand, heterochromatic RNA reduces the mobility and spreading of the fission yeast HP1 protein in heterochromatic domains (Keller et al., 2012; Muchardt et al., 2002; Stunnenberg et al., 2015). From an evolutionary perspective, the properties of co-localization, abundance, and sequence preference of L1 and HP1α might have co-evolved to ensure regulated formation of L1-enriched heterochromatin in organizing the nucleus. L1 RNA binds NCL, and its depletion causes relocation of *DUX* genes from the perinucleolar heterochromatin to the nucleoplasm (Percharde et al., 2018). Abnormal activation or silencing of L1 in mouse zygotes led to altered chromatin accessibility and compaction, which failed to be rescued by synthetic L1 RNA (Jachowicz et al., 2017). Together with previous reports, our results suggest a model where L1 RNA acts in *cis* to anchor L1 repeat-enriched chromosomal segments to the nucleolar and nuclear peripheries. This scaffold function is reminiscent of reported roles for several chromatin-bound noncoding RNAs in organizing subnuclear domains. For example, *Xist* RNA binds the lamin B receptor in the inner nuclear membrane to initiate and maintain the silencing of the inactive X chromosome (McHugh et al., 2015). Transcription at major satellite repeats precedes chromocenter formation, and satellite transcripts help to recruit the H3K9me3 methyltransferase SUV39H to centromeric DNA sequences (Probst et al., 2010; Velazquez Camacho et al., 2017). However, the exact mechanism underlying the function of L1 RNA remains to be fully elucidated by future studies.

In summary, our study starts to unravel the fundamental principle of 3D genome organization, in which the distribution of L1 and B1 transposable elements provides the blueprint for chromatin folding. As discovered by Anfinsen in the late 1950s, the central pillar of protein science tells us that the amino acid sequence of a protein determines its structure and function (Anfinsen, 1972; Anfinsen et al., 1954). Analogously, we propose that the primary DNA sequences determine how the genome folds and functions. We believe that genome folding occurs autonomously, through a process that is driven by homotypic clustering of regions containing L1 or B1 repeat sequences, and is further facilitated by transcription and transcripts produced at these repeat elements. Structural information embedded in L1 and SINE repeats may be universally recognizable, thus contributing to the high degree of stability and conservation in compartmental and TAD organization that is observed across mammalian species and cell types. We want to note that L1 and B1/Alu compartmental domains represent a structural and functional ground state of chromatin organization, on which subsequent regulatory features, such as dynamic enhancer-promoter interactions, are overlaid. Nevertheless, the same principle of homotypic clustering, phase separation and spatial segregation of chromosomal segments may be reiterated at different genomic scales, consequently folding the genome.

## ACKNOWLEDGMENTS

We thank the Shen Laboratory members for insightful discussions, and we thank Dr. Hongxia Lv at the core imaging facility of the School of Life Sciences, Peking University, for imaging support. This work was supported in part by the National Basic Research Program of China (2018YFA0107604, 2017YFA0504204 to X.S.); the National Natural Science Foundation of China (31630095, 31471219 to X.S.; 21573013, 21825401 to Y.S.); the National Key R&D Program of China (2017YFA0505300 to Y.S.); and the Center for Life Sciences (CLS) at Tsinghua University (to X.S).

## AUTHOR CONTRIBUTIONS

X.S. conceived of and supervised the study. X.S., Y.S., J.Y.L. and L.C. designed the experiments. J.Y.L. performed bioinformatics analysis. L.C. conducted all imaging experiments with help from Y.Y., G.Z., T.W., X.H., and J.Z. T.L. performed phase separation and RNA pull-down experiments. K.Z., Q.P. and P.Y. prepared the Hi-C library. T.W., W.L. and P.W. did embryo injections. P.L. and L.W. provide HP1α proteins. M.P and M.R.-S. contributed to early experiments of L1 RNA knockdown. W.X., J.N., H.Z., X.B., W.S., Z.W. and Y.H. contributed technical assistance/suggestions. X.S., J.Y.L. and L.C. wrote the manuscript with input from all authors.

## REFERENCES

Anfinsen, C.B. (1972). The formation and stabilization of protein structure. Biochem J 128, 737–749.

Anfinsen, C.B., Redfield, R.R., Choate, W.L., Page, J., and Carroll, W.R. (1954). Studies on the gross structure, cross-linkages, and terminal sequences in ribonuclease. J Biol Chem 207, 201–210.

Annunziato, A.T. (2008). DNA Packaging: Nucleosomes and Chromatin. Nature Education 1(1):26.

Bergeron-Sandoval, L.P., Safaee, N., and Michnick, S.W. (2016). Mechanisms and Consequences of Macromolecular Phase Separation. Cell 165, 1067–1079.

Biemont, C. (2010). A Brief History of the Status of Transposable Elements: From Junk DNA to Major Players in Evolution. Genetics 186, 1085–1093.

Bintu, B., Mateo, L.J., Su, J.H., Sinnott-Armstrong, N.A., Parker, M., Kinrot, S., Yamaya, K., Boettiger, A.N., and Zhuang, X. (2018). Super-resolution chromatin tracing reveals domains and cooperative interactions in single cells. Science 362.

Boettiger, A.N., Bintu, B., Moffitt, J.R., Wang, S., Beliveau, B.J., Fudenberg, G., Imakaev, M., Mirny, L.A., Wu, C.T., and Zhuang, X. (2016). Super-resolution imaging reveals distinct chromatin folding for different epigenetic states. Nature 529, 418–422.

Bonev, B., and Cavalli, G. (2016). Organization and function of the 3D genome. Nat Rev Genet 17, 661–678.

Bourque, G., Leong, B., Vega, V.B., Chen, X., Lee, Y.L., Srinivasan, K.G., Chew, J.L., Ruan, Y., Wei, C.L., Ng, H.H., et al. (2008). Evolution of the mammalian transcription factor binding repertoire via transposable elements. Genome Res 18, 1752–1762.

Bouwman, B.A., and de Laat, W. (2015). Getting the genome in shape: the formation of loops, domains and compartments. Genome Biol 16, 154.

Buchwalter, A., Kaneshiro, J.M., and Hetzer, M.W. (2019). Coaching from the sidelines: the nuclear periphery in genome regulation. Nature Reviews Genetics 20, 39–50.

Chen, C., Ara, T., and Gautheret, D. (2009). Using Alu elements as polyadenylation sites: A case of retroposon exaptation. Mol Biol Evol 26, 327–334.

Chong, S., Dugast-Darzacq, C., Liu, Z., Dong, P., Dailey, G.M., Cattoglio, C., Heckert, A., Banala, S., Lavis, L., Darzacq, X., et al. (2018). Imaging dynamic and selective low-complexity domain interactions that control gene transcription. Science 361.

Chuong, E.B., Elde, N.C., and Feschotte, C. (2016). Regulatory evolution of innate immunity through co-option of endogenous retroviruses. Science 351, 1083–1087.

Cook, P.R. (2002). Predicting three-dimensional genome structure from transcriptional activity. Nat Genet 32, 347–352.

Cook, P.R., and Marenduzzo, D. (2018). Transcription-driven genome organization: a model for chromosome structure and the regulation of gene expression tested through simulations. Nucleic Acids Res 46, 9895–9906.

Deininger, P. (2011). Alu elements: know the SINEs. Genome Biol 12, 236.

Dekker, J., Belmont, A.S., Guttman, M., Leshyk, V.O., Lis, J.T., Lomvardas, S., Mirny, L.A., O’Shea, C.C., Park, P.J., Ren, B., et al. (2017). The 4D nucleome project. Nature 549, 219–226.

Dekker, J., Marti-Renom, M.A., and Mirny, L.A. (2013). Exploring the three-dimensional organization of genomes: interpreting chromatin interaction data. Nat Rev Genet 14, 390–403.

Dekker, J., Rippe, K., Dekker, M., and Kleckner, N. (2002). Capturing chromosome conformation. Science 295, 1306–1311.

Dixon, J.R., Jung, I., Selvaraj, S., Shen, Y., Antosiewicz-Bourget, J.E., Lee, A.Y., Ye, Z., Kim, A., Rajagopal, N., Xie, W., et al. (2015). Chromatin architecture reorganization during stem cell differentiation. Nature 518, 331–336.

Dixon, J.R., Selvaraj, S., Yue, F., Kim, A., Li, Y., Shen, Y., Hu, M., Liu, J.S., and Ren, B. (2012). Topological domains in mammalian genomes identified by analysis of chromatin interactions. Nature 485, 376–380.

Dominissini, D., Moshitch-Moshkovitz, S., Amariglio, N., and Rechavi, G. (2011). Adenosine-to-inosine RNA editing meets cancer. Carcinogenesis 32, 1569–1577.

Dowen, J.M., Fan, Z.P., Hnisz, D., Ren, G., Abraham, B.J., Zhang, L.N., Weintraub, A.S., Schujiers, J., Lee, T.I., Zhao, K., et al. (2014). Control of cell identity genes occurs in insulated neighborhoods in mammalian chromosomes. Cell 159, 374–387.

Du, Z., Zheng, H., Huang, B., Ma, R., Wu, J., Zhang, X., He, J., Xiang, Y., Wang, Q., Li, Y., et al. (2017). Allelic reprogramming of 3D chromatin architecture during early mammalian development. Nature 547, 232–235.

Durruthy-Durruthy, J., Sebastiano, V., Wossidlo, M., Cepeda, D., Cui, J., Grow, E.J., Davila, J., Mall, M., Wong, W.H., Wysocka, J., et al. (2016). The primate-specific noncoding RNA HPAT5 regulates pluripotency during human preimplantation development and nuclear reprogramming. Nat Genet 48, 44–52.

Fadloun, A., Le Gras, S., Jost, B., Ziegler-Birling, C., Takahashi, H., Gorab, E., Carninci, P., and Torres-Padilla, M.E. (2013). Chromatin signatures and retrotransposon profiling in mouse embryos reveal regulation of LINE-1 by RNA. Nat Struct Mol Biol 20, 332–338.

Falahati, H., Pelham-Webb, B., Blythe, S., and Wieschaus, E. (2016). Nucleation by rRNA Dictates the Precision of Nucleolus Assembly. Curr Biol 26, 277–285.

Falk, M., Feodorova, Y., Naumova, N., Imakaev, M., Lajoie, B.R., Leonhardt, H., Joffe, B., Dekker, J., Fudenberg, G., Solovei, I., et al. (2019). Heterochromatin drives compartmentalization of inverted and conventional nuclei. Nature 570, 395–399.

Flyamer, I.M., Gassler, J., Imakaev, M., Brandao, H.B., Ulianov, S.V., Abdennur, N., Razin, S.V., Mirny, L.A., and Tachibana-Konwalski, K. (2017). Single-nucleus Hi-C reveals unique chromatin reorganization at oocyte-to-zygote transition. Nature 544, 110–114.

Fraser, J., Ferrai, C., Chiariello, A.M., Schueler, M., Rito, T., Laudanno, G., Barbieri, M., Moore, B.L., Kraemer, D.C., Aitken, S., et al. (2015). Hierarchical folding and reorganization of chromosomes are linked to transcriptional changes in cellular differentiation. Mol Syst Biol 11, 852.

Gassler, J., Brandao, H.B., Imakaev, M., Flyamer, I.M., Ladstatter, S., Bickmore, W.A., Peters, J.M., Mirny, L.A., and Tachibana, K. (2017). A mechanism of cohesin-dependent loop extrusion organizes zygotic genome architecture. Embo J 36, 3600–3618.

Gibcus, J.H., and Dekker, J. (2013). The hierarchy of the 3D genome. Mol Cell 49, 773–782.

Grow, E.J., Flynn, R.A., Chavez, S.L., Bayless, N.L., Wossidlo, M., Wesche, D.J., Martin, L., Ware, C.B., Blish, C.A., Chang, H.Y., et al. (2015). Intrinsic retroviral reactivation in human preimplantation embryos and pluripotent cells. Nature 522, 221–225.

Guelen, L., Pagie, L., Brasset, E., Meuleman, W., Faza, M.B., Talhout, W., Eussen, B.H., de Klein, A., Wessels, L., de Laat, W., et al. (2008). Domain organization of human chromosomes revealed by mapping of nuclear lamina interactions. Nature 453, 948–951.

Haarhuis, J.H.I., van der Weide, R.H., Blomen, V.A., Yanez-Cuna, J.O., Amendola, M., van Ruiten, M.S., Krijger, P.H.L., Teunissen, H., Medema, R.H., van Steensel, B., et al. (2017). The Cohesin Release Factor WAPL Restricts Chromatin Loop Extension. Cell 169, 693–707 e614.

Houda Belaghzal, T.B., Andrew D. Stephens, Denis L. Lafontaine, Sergey V. Venev, Zhiping Weng, John F. Marko, Job Dekker (2019). Compartment-dependent chromatin interaction dynamics revealed by liquid chromatin Hi-C. bioRxiv.

Jachowicz, J.W., Bing, X.Y., Pontabry, J., Boskovic, A., Rando, O.J., and Torres-Padilla, M.E. (2017). LINE-1 activation after fertilization regulates global chromatin accessibility in the early mouse embryo. Nature Genetics 49, 1502–1510.

Jost, D., Carrivain, P., Cavalli, G., and Vaillant, C. (2014). Modeling epigenome folding: formation and dynamics of topologically associated chromatin domains. Nucleic Acids Res 42, 9553–9561.

Jurka, J., Kohany, O., Pavlicek, A., Kapitonov, V.V., and Jurka, M.V. (2004). Duplication, coclustering, and selection of human Alu retrotransposons. Proc Natl Acad Sci U S A 101, 1268–1272.

Kadauke, S., and Blobel, G.A. (2009). Chromatin loops in gene regulation. Biochim Biophys Acta 1789, 17–25.

Ke, Y., Xu, Y., Chen, X., Feng, S., Liu, Z., Sun, Y., Yao, X., Li, F., Zhu, W., Gao, L., et al. (2017). 3D Chromatin Structures of Mature Gametes and Structural Reprogramming during Mammalian Embryogenesis. Cell 170, 367–381 e320.

Keller, C., Adaixo, R., Stunnenberg, R., Woolcock, K.J., Hiller, S., and Buhler, M. (2012). HP1(Swi6) mediates the recognition and destruction of heterochromatic RNA transcripts. Mol Cell 47, 215–227.

Korenberg, J.R., and Rykowski, M.C. (1988). Human genome organization: Alu, lines, and the molecular structure of metaphase chromosome bands. Cell 53, 391–400.

Lander, E.S., Linton, L.M., Birren, B., Nusbaum, C., Zody, M.C., Baldwin, J., Devon, K., Dewar, K., Doyle, M., FitzHugh, W., et al. (2001). Initial sequencing and analysis of the human genome. Nature 409, 860–921.

Larson, A.G., Elnatan, D., Keenen, M.M., Trnka, M.J., Johnston, J.B., Burlingame, A.L., Agard, D.A., Redding, S., and Narlikar, G.J. (2017). Liquid droplet formation by HP1alpha suggests a role for phase separation in heterochromatin. Nature 547, 236–240.

Li, P., Banjade, S., Cheng, H.C., Kim, S., Chen, B., Guo, L., Llaguno, M., Hollingsworth, J.V., King, D.S., Banani, S.F., et al. (2012). Phase transitions in the assembly of multivalent signalling proteins. Nature 483, 336–340.

Lieberman-Aiden, E., van Berkum, N.L., Williams, L., Imakaev, M., Ragoczy, T., Telling, A., Amit, I., Lajoie, B.R., Sabo, P.J., Dorschner, M.O., et al. (2009). Comprehensive mapping of long-range interactions reveals folding principles of the human genome. Science 326, 289–293.

Liu, N., Lee, C.H., Swigut, T., Grow, E., Gu, B., Bassik, M.C., and Wysocka, J. (2018). Selective silencing of euchromatic L1s revealed by genome-wide screens for L1 regulators. Nature 553, 228–232.

Lubelsky, Y., and Ulitsky, I. (2018). Sequences enriched in Alu repeats drive nuclear localization of long RNAs in human cells. Nature 555, 107–111.

Lynch, V.J., Leclerc, R.D., May, G., and Wagner, G.P. (2011). Transposon-mediated rewiring of gene regulatory networks contributed to the evolution of pregnancy in mammals. Nat Genet 43, 1154–1159.

Mandal, P.K., and Kazazian, H.H., Jr. (2008). SnapShot: Vertebrate transposons. Cell 135, 192–192 e191.

Mateos-Langerak, J., Brink, M.C., Luijsterburg, M.S., van der Kraan, I., van Driel, R., and Verschure, P.J. (2007). Pericentromeric heterochromatin domains are maintained without accumulation of HP1. Molecular Biology of the Cell 18, 1464–1471.

McHugh, C.A., Chen, C.K., Chow, A., Surka, C.F., Tran, C., McDonel, P., Pandya-Jones, A., Blanco, M., Burghard, C., Moradian, A., et al. (2015). The Xist lncRNA interacts directly with SHARP to silence transcription through HDAC3. Nature 521, 232–236.

Meuleman, W., Peric-Hupkes, D., Kind, J., Beaudry, J.B., Pagie, L., Kellis, M., Reinders, M., Wessels, L., and van Steensel, B. (2013). Constitutive nuclear lamina-genome interactions are highly conserved and associated with A/T-rich sequence. Genome Res 23, 270–280.

Mirny, L.A. (2011). The fractal globule as a model of chromatin architecture in the cell. Chromosome Res 19, 37–51.

Misteli, T. (2007). Beyond the sequence: cellular organization of genome function. Cell 128, 787–800.

Mouse Genome Sequencing, C., Waterston, R.H., Lindblad-Toh, K., Birney, E., Rogers, J., Abril, J.F., Agarwal, P., Agarwala, R., Ainscough, R., Alexandersson, M., et al. (2002). Initial sequencing and comparative analysis of the mouse genome. Nature 420, 520–562.

Muchardt, C., Guilleme, M., Seeler, J.S., Trouche, D., Dejean, A., and Yaniv, M. (2002). Coordinated methyl and RNA binding is required for heterochromatin localization of mammalian HP1 alpha. Embo Reports 3, 975–981.

Nagano, T., Lubling, Y., Vaarnai, C., Dudley, C., Leung, W., Baran, Y., Cohen, N.M., Wingett, S., Fraser, P., and Tanay, A. (2017). Cell-cycle dynamics of chromosomal organization at single-cell resolution. Nature 547, 61–67.

Naumova, N., Imakaev, M., Fudenberg, G., Zhan, Y., Lajoie, B.R., Mirny, L.A., and Dekker, J. (2013). Organization of the mitotic chromosome. Science 342, 948–953.

Nora, E.P., Goloborodko, A., Valton, A.L., Gibcus, J.H., Uebersohn, A., Abdennur, N., Dekker, J., Mirny, L.A., and Bruneau, B.G. (2017). Targeted Degradation of CTCF Decouples Local Insulation of Chromosome Domains from Genomic Compartmentalization. Cell 169, 930–944.

Nuebler, J., Fudenberg, G., Imakaev, M., Abdennur, N., and Mirny, L.A. (2018). Chromatin organization by an interplay of loop extrusion and compartmental segregation. Proc Natl Acad Sci U S A 115, E6697–E6706.

Orgel, L.E., and Crick, F.H. (1980). Selfish DNA: the ultimate parasite. Nature 284, 604–607.

Percharde, M., Lin, C.J., Yin, Y., Guan, J., Peixoto, G.A., Bulut-Karslioglu, A., Biechele, S., Huang, B., Shen, X., and Ramalho-Santos, M. (2018). A LINE1-Nucleolin Partnership Regulates Early Development and ESC Identity. Cell 174, 391–405 e319.

Polak, P., and Domany, E. (2006). Alu elements contain many binding sites for transcription factors and may play a role in regulation of developmental processes. Bmc Genomics 7, 133.

Probst, A.V., Okamoto, I., Casanova, M., El Marjou, F., Le Baccon, P., and Almouzni, G. (2010). A strand-specific burst in transcription of pericentric satellites is required for chromocenter formation and early mouse development. Dev Cell 19, 625–638.

Pueschel, R., Coraggio, F., and Meister, P. (2016). From single genes to entire genomes: the search for a function of nuclear organization. Development 143, 910–923.

Qiu, J.J., Zhang, W.W., Wu, Z.L., Wang, Y.H., Qian, M., and Li, Y.P. (2003). Delay of ZGA initiation occurred in 2-cell blocked mouse embryos. Cell Res 13, 179–185.

Quinodoz, S.A., Ollikainen, N., Tabak, B., Palla, A., Schmidt, J.M., Detmar, E., Lai, M.M., Shishkin, A.A., Bhat, P., Takei, Y., et al. (2018). Higher-Order Inter-chromosomal Hubs Shape 3D Genome Organization in the Nucleus. Cell 174, 744–757 e724.

Rao, S.S., Huntley, M.H., Durand, N.C., Stamenova, E.K., Bochkov, I.D., Robinson, J.T., Sanborn, A.L., Machol, I., Omer, A.D., Lander, E.S., et al. (2014). A 3D map of the human genome at kilobase resolution reveals principles of chromatin looping. Cell 159, 1665–1680.

Rao, S.S.P., Huang, S.C., St Hilaire, B.G., Engreitz, J.M., Perez, E.M., Kieffer-Kwon, K.R., Sanborn, A.L., Johnstone, S.E., Bascom, G.D., Bochkov, I.D., et al. (2017). Cohesin Loss Eliminates All Loop Domains. Cell 171, 305–320.

Rebollo, R., Romanish, M.T., and Mager, D.L. (2012). Transposable Elements: An Abundant and Natural Source of Regulatory Sequences for Host Genes. Annual Review of Genetics, Vol 46 *46*, 21–42.

Rivera-Mulia, J.C., and Gilbert, D.M. (2016). Replication timing and transcriptional control: beyond cause and effect-part III. Curr Opin Cell Biol 40, 168–178.

Rothstein, J.L., Johnson, D., Deloia, J.A., Skowronski, J., Solter, D., and Knowles, B. (1992). Gene-Expression during Preimplantation Mouse Development. Gene Dev 6, 1190–1201.

Rowley, M.J., and Corces, V.G. (2018). Organizational principles of 3D genome architecture. Nat Rev Genet 19, 789–800.

Rowley, M.J., Nichols, M.H., Lyu, X.W., Ando-Kuri, M., Rivera, I.S.M., Hermetz, K., Wang, P., Ruan, Y.J., and Corces, V.G. (2017). Evolutionarily Conserved Principles Predict 3D Chromatin Organization. Mol Cell 67, 837–852.

Schwarzer, W., Abdennur, N., Goloborodko, A., Pekowska, A., Fudenberg, G., Loe-Mie, Y., Fonseca, N.A., Huber, W., C, H.H., Mirny, L., et al. (2017). Two independent modes of chromatin organization revealed by cohesin removal. Nature 551, 51–56.

Solovei, I., Thanisch, K., and Feodorova, Y. (2016). How to rule the nucleus: divide et impera. Current Opinion in Cell Biology 40, 47–59.

Splinter, E., Heath, H., Kooren, J., Palstra, R.J., Klous, P., Grosveld, F., Galjart, N., and de Laat, W. (2006). CTCF mediates long-range chromatin looping and local histone modification in the beta-globin locus. Genes Dev 20, 2349–2354.

Stadhouders, R., Vidal, E., Serra, F., Di Stefano, B., Le Dily, F., Quilez, J., Gomez, A., Collombet, S., Berenguer, C., Cuartero, Y., et al. (2018). Transcription factors orchestrate dynamic interplay between genome topology and gene regulation during cell reprogramming. Nature Genetics 50, 238–249.

Stevens, T.J., Lando, D., Basu, S., Atkinson, L.P., Cao, Y., Lee, S.F., Leeb, M., Wohlfahrt, K.J., Boucher, W., O’Shaughnessy-Kirwan, A., et al. (2017). 3D structures of individual mammalian genomes studied by single-cell Hi-C. Nature 544, 59–64.

Strom, A.R., Emelyanov, A.V., Mir, M., Fyodorov, D.V., Darzacq, X., and Karpen, G.H. (2017). Phase separation drives heterochromatin domain formation. Nature 547, 241–245.

Stunnenberg, R., Kulasegaran-Shylini, R., Keller, C., Kirschmann, M.A., Gelman, L., and Buhler, M. (2015). H3K9 methylation extends across natural boundaries of heterochromatin in the absence of an HP1 protein. Embo J 34, 2789–2803.

Sundaram, V., and Wang, T. (2018). Transposable Element Mediated Innovation in Gene Regulatory Landscapes of Cells: Re-Visiting the “Gene-Battery” Model. Bioessays 40.

Tang, S.J. (2017). Potential Role of Phase Separation of Repetitive DNA in Chromosomal Organization. Genes (Basel) 8.

Taylor, M.S., LaCava, J., Mita, P., Molloy, K.R., Huang, C.R., Li, D., Adney, E.M., Jiang, H., Burns, K.H., Chait, B.T., et al. (2013). Affinity proteomics reveals human host factors implicated in discrete stages of LINE-1 retrotransposition. Cell 155, 1034–1048.

Tjong, H., Li, W.Y., Kalhor, R., Dai, C., Hao, S.L., Gong, K., Zhou, Y.G., Li, H.C., Zhou, X.J., Le Gros, M.A., et al. (2016). Population-based 3D genome structure analysis reveals driving forces in spatial genome organization. P Natl Acad Sci USA 113, E1663–E1672.

van de Werken, H.J.G., Haan, J.C., Feodorova, Y., Bijos, D., Weuts, A., Theunis, K., Holwerda, S.J.B., Meuleman, W., Pagie, L., Thanisch, K., et al. (2017). Small chromosomal regions position themselves autonomously according to their chromatin class. Genome Res 27, 922–933.

van Steensel, B., and Belmont, A.S. (2017). Lamina-Associated Domains: Links with Chromosome Architecture, Heterochromatin, and Gene Repression. Cell 169, 780–791.

Velazquez Camacho, O., Galan, C., Swist-Rosowska, K., Ching, R., Gamalinda, M., Karabiber, F., De La Rosa-Velazquez, I., Engist, B., Koschorz, B., Shukeir, N., et al. (2017). Major satellite repeat RNA stabilize heterochromatin retention of Suv39h enzymes by RNA-nucleosome association and RNA:DNA hybrid formation. Elife 6.

Wang, J., Xie, G., Singh, M., Ghanbarian, A.T., Rasko, T., Szvetnik, A., Cai, H., Besser, D., Prigione, A., Fuchs, N.V., et al. (2014). Primate-specific endogenous retrovirus-driven transcription defines naive-like stem cells. Nature 516, 405–409.

Wang, S., Su, J.H., Beliveau, B.J., Bintu, B., Moffitt, J.R., Wu, C.T., and Zhuang, X. (2016). Spatial organization of chromatin domains and compartments in single chromosomes. Science 353, 598–602.

Wijchers, P.J., Geeven, G., Eyres, M., Bergsma, A.J., Janssen, M., Verstegen, M., Zhu, Y., Schell, Y., Vermeulen, C., de Wit, E., et al. (2015). Characterization and dynamics of pericentromere-associated domains in mice. Genome Res 25, 958–969.

Wutz, G., Várnai, C., Nagasaka, K., Cisneros, D.A., Stocsits, R.R., Tang, W., Schoenfelder, S., Jessberger, G., Muhar, M., Hossain, M.J., et al. (2017). Topologically associating domains and chromatin loops depend on cohesin and are regulated by CTCF, WAPL, and PDS5 proteins. The EMBO journal 36, 3573–3599.

Xie, Y., Rosser, J.M., Thompson, T.L., Boeke, J.D., and An, W. (2010). Characterization of L1 retrotransposition with high-throughput dual-luciferase assays. Nucleic acids research 39, e16–e16.

Zane, L., Sharma, V., and Misteli, T. (2014). Common features of chromatin in aging and cancer: cause or coincidence? Trends Cell Biol 24, 686–694.

Zullo, J.M., Demarco, I.A., Pique-Regi, R., Gaffney, D.J., Epstein, C.B., Spooner, C.J., Luperchio, T.R., Bernstein, B.E., Pritchard, J.K., Reddy, K.L., et al. (2012). DNA sequence-dependent compartmentalization and silencing of chromatin at the nuclear lamina. Cell 149, 1474–1487.

Zwicker, D., Decker, M., Jaensch, S., Hyman, A.A., and Julicher, F. (2014). Centrosomes are autocatalytic droplets of pericentriolar material organized by centrioles. P Natl Acad Sci USA 111, E2636–E2645.

